# A 30-Color Full-Spectrum Flow Cytometry Panel to Characterize the Immune Cell Landscape in Spleen and Tumors within a Syngeneic MC-38 Murine Colon Carcinoma Model

**DOI:** 10.1101/2023.02.24.529779

**Authors:** Gabriel DeNiro, Kathryn Que, Soo Min Koo, Jeong Kim, Bridget Schneider, Anandaroop Mukhopadhyay, Anandi Sawant, Tuan Andrew Nguyen

## Abstract

**Purpose and Appropriate Sample Types:** This panel was designed to characterize the immune cell landscape in the mouse tumor micro-environment as well as mouse lymphoid tissues (e.g., spleen). Previous Optimized Multicolor Immunofluorescence Panels (OMIP) with conventional cytometry were examples of high-quality fluorescent-based flow cytometry panels to characterize either the T-cell compartment or the myeloid compartment [1, 2]. The advent of spectral cytometry has enabled sufficient markers to be included in a panel to comprehensively characterize the immune cell landscape including both the T-cell and the myeloid cell compartments. In this body of work, we demonstrated that we could measure the frequency and characterize the functional status of CD4 T cells, CD8 T cells, regulatory T cells, NK cells, B cells, macrophages, granulocytes, monocytes, & dendritic cells. This panel is especially useful for understanding the immune landscape in “cold” preclinical tumor models with very low immune cell infiltration, and for investigating how therapeutic treatments may modulate the immune landscape.

## Background

Immunotherapy has revolutionized the landscape of cancer care treatment, and in some patients, it has resulted in a long-lasting anti-tumor response. The success of Bristol-Myers Squibb’s ipilimumab (anti-CTLA4 antibody), nivolumab (anti-PD1 antibody), and relatlimab (anti-LAG3 antibody) relies on inducing an immune response of pre-existing T cells whose activity was limited due to immune checkpoints [3]. Unfortunately, many patients still do not respond to treatments [4], prompting scientists to study the mechanism of resistance [5] and identify novel therapeutics and combination strategies that can enhance the response rate [6].

In the research and development for the next-generation of immunotherapy, scientists typically rely on preclinical syngeneic mouse-tumor models, in which immortalized mouse cancer cell lines are engrafted into inbred mice with a matching genetic background and a competent immune system, to demonstrate drug efficacy. In these preclinical studies, flow cytometry is a quintessential technique. Single cell suspensions from tumor and spleen are examined by flow cytometry to characterize the immune cell landscape and drug pharmacodynamics. Because drug development is an iterative process, insights gained from flow cytometry data help inform on the drug’s mechanism of action, drug lead optimization, combination strategy with other drugs, and dosing strategy [7–9].

Historically, scientists using the conventional BD LSR Fortessa have been limited to having only up to 18 colors in each panel. As a result, to comprehensively profile the immune cell landscape, scientists typically need at least two panels: a lymphoid-focused panel or a myeloid-focused panel. This was previously echoed in OMIP-31 and OMIP-32 multicolor panels [1, 2]. On the other hand, tumor samples are often limited, and as a result, there are only enough cells to run one panel. While there was effort to establish a high dimensional panel to broadly characterize the immune cell landscape by mass cytometry [10], the low sample throughput rate (1000 cells/s) and the lack of a 96-well plate reader [11] have limited the adoption of mass cytometry in these preclinical studies. Despite the popularity of syngeneic mouse tumor models in preclinical research, there has been no previously published high-dimensional fluorescent OMIP panel to characterize the mouse tumor immune landscape beyond 16 fluorescent parameters.

In this paper, we propose a 30-color flow cytometry approach to comprehensively characterize the immune cell landscape, relying on the spectral capability of the BD FACS Symphony A5 SE. To achieve this, we carefully selected 30 reagents to measure the frequency of canonical immune cell populations and characterize their phenotype and functionality. The choices of antibody clones, fluorochromes, markers were driven by their compatibility with the tumor dissociation enzymes, fluorochrome brightness, antigen density, instrument configuration, and instrument spillover spread matrix (Figure S1-S5). We have also provided a detailed tissue processing protocol (Figure S6) and flow cytometry staining protocol (Figure S7).

A representative gating strategy to delineate immune cell subsets is detailed in Figure 1A. Briefly, dissociated tumor cells were first gated on CD45+ to capture the immune cells, followed by viability gating (Live/Dead Blue negative) and singlet gating. Subsequently, we plotted CD11b against NK1.1 to identify CD11b+ myeloid cells, NK1.1+ NK cells, and other CD11b- NK1.1- immune cells.

**Figure 1.**
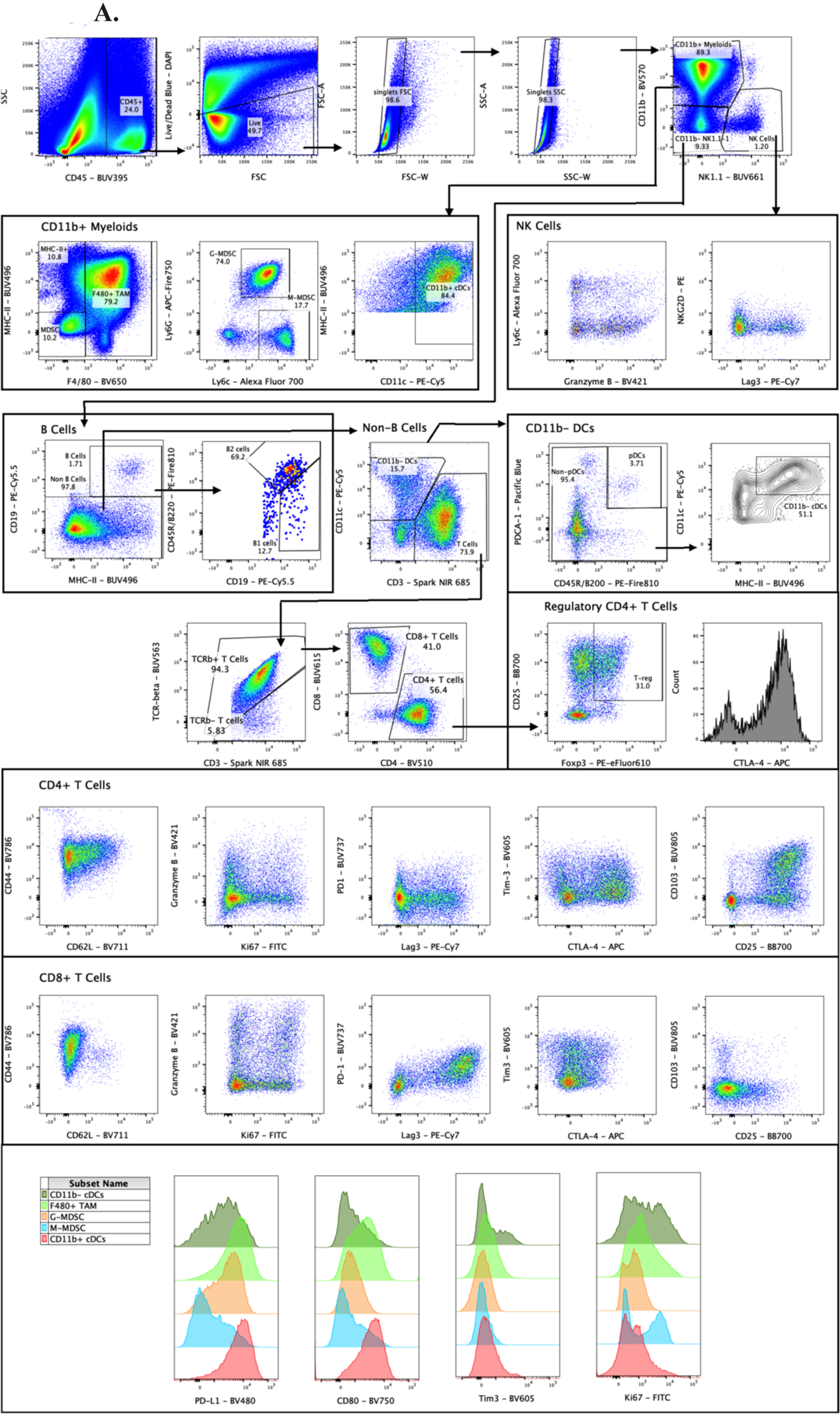
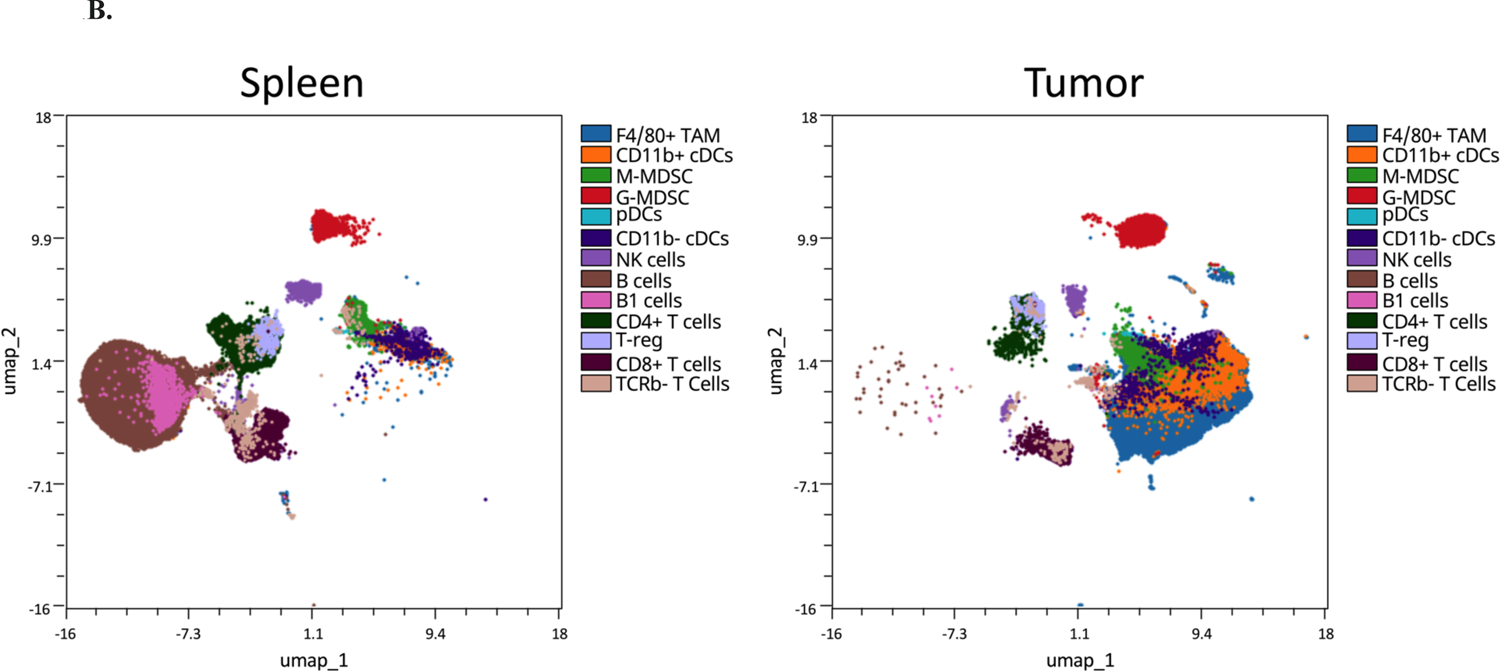
Validation of the 30-color panel with syngeneic mouse MC38 tumors. **A) An example 2×2 gating strategy of the sample.** After gating on CD45+, live, FSC singlets, SSC singlets, cells were gated on CD11b and NK1.1 to separate CD11b+ myeloid cells, NK cells, and CD11b-/NK1.1- cells. The CD11b+ myeloid cells were further divided in Tumor-Associated Macrophages, CD11b+ dendritic cells, G-MDSC, M-MDSC based on the expressions of MHC-II, F4/80, CD11c, Ly6C, and Ly6G. The CD11b-/NK1.1- fractions were divided into B cells (CD19+ / MHC-II+) and non-B cells (CD19-). The non-B cells was further fractionated into T cells and CD11b- dendritic cells based on expression CD3 and CD11c. Out of the CD11b- dendritic cells, pDCs were identified based on CD45R/B220 and PDCA-1 expression, and CD11b- conventional dendritic cells were identified based on the high expression of CD11c and MHC-II+. Among the CD3+ T cells, TCR-beta was used to classify T cells into TCR-beta+ T cells and TCR-beta-T cells. Helper CD4 and Cytotoxic CD8 T cells were identified from the TCR-beta+ T cells gate. Regulatory CD4+ T cells were identified from the CD4+ T cell gate using CD25 and Foxp3 (CD25+/Foxp3+). **B) UMAP dimensionality reduction analysis of the live CD45+ immune cells from the spleen and tumor.** A legend to depict each population was included.

CD11b+ myeloid was further divided into granulocytic myeloid-derived suppressive cells (G-MDSC) (CD11b+ F4/80- MHC-II- Ly6G+ Ly6Cmed), monocytic myeloid-derived suppressive cells (M-MDSC) (CD11b+ F4/80- MHC-II- Ly6G- Ly6Chi) [12, 13], tumor-associated macrophages (TAM) (CD11b+ F4/80+) and CD11b+ conventional dendritic cells (CD11b+ cDCs) (CD11b+ F4/80- MHC-II+ CD11c+) [14].

The NK1.1+ NK cells can be further characterized by their expression of Ly6c, Granzyme B, LAG-3, and NKG2D. Ly6c is a marker for a subset of mature and resting NK cells with a lower level of Granzyme B production, compared to Ly6c-NK cells [15]. NKG2D is an activating receptors that can be expressed on NK cells and CD8+ T cells [16], and an immunotherapy to upregulate NKG2D expression on immune cells may contribute to a robust anti-tumor response [17, 18]. LAG-3+ NK cells represent a subset with impaired cytokine production [19] and reduced target cell killing activity [20].

CD11b- NK1.1- immune cells were divided into B cells (CD19+ MHC-II+), T cells (CD19-, CD3+, CD11c+/-), plasmacytoid dendritic cells (pDCs) (CD19-, CD3-, CD11c+, PDCA-1+, CD45/B220+) [21], and CD11b- conventional dendritic cells (CD11b- cDCs) (CD19-, CD3-, CD11b-, CD11c+, MHC-II+) [14]. Although not included in this example gating strategy, CD103, CD4, CD8 can be used to further characterize CD11b+ conventional dendritic cell and CD11b- conventional dendritic cells [22]. Among the B cells, they can be further divided into B1 cells (CD19+ CD45R/B220 low) and B2 cells (CD19+CD45R/B220 hi) [23]. However, the role of B1 cells and B2 cells in anti-tumor immunity is still unclear [24].

Among the T cells, it was divided into TCR-beta+ and TCR-beta- subsets. TCR-beta positive refers to all alpha/beta T cells and represents the main pool of T cells [25]; on the other hands, TCR-beta negative corresponds to gamma/delta T cells - a minor subset of T cells that may regulate cancer development and progression [26]. The TCR-Beta+ subset is further divided into T-helper and T-cytotoxic cells based on the expression of CD4 and CD8, respectively. CD4+ T-regulatory cells are identified as CD25+/Foxp3+ double positive. Other markers such as CD44, CD62L, Granzyme B, Ki-67, CD103, PD-1, LAG-3, Tim-3, CTLA-4 are included to characterize the differentiation and functionality of T cells. In mice, CD44 and CD62L are classical markers to characterize naïve (CD44-, CD62L+), central memory (CD44+CD62L+), effector memory (CD44+CD62L-) T cells [1]. The ratio of central memory over effector T cells can be a predictive biomarker of immunotherapy efficacy [27]. Granzyme B was previously shown to be an early predictor of therapeutic efficacy for checkpoint inhibitor regimens in preclinical settings [28]. Detection of Ki-67 among PD-1+CD8+ T cells was demonstrated as a biomarker to predict the response and prognosis to checkpoint inhibitor regimens in clinical trials [29, 30]. The infiltration of intratumoral CD103+CD8+ T cell can predict response to checkpoint inhibitor [31]. PD-1, CTLA-4, LAG-3 and Tim-3 are well-described checkpoint markers that define T-cell exhaustion and primary resistance mechanism to immunotherapy [32–35]. Among these checkpoint markers, therapeutics targeting PD-1, CTLA-4, and LAG-3 from Bristol Myers Squibb have been approved based on demonstrated clinical benefits in cancer patients [36–38]. The inclusion of these checkpoint markers in the flow cytometry immunophenotyping panel not only describes the exhaustion status of T-cell, but also informs on potential combo-therapy strategy of drug candidates with these existing approved therapeutics.

Lastly, among the myeloid cell subsets, we can measure their expression of PD-L1, CD80, Tim-3, and Ki-67. PD-L1 expression on intratumoral myeloid cells is known to regulate anti-tumor immune response [39–41]. CD80 is one of the costimulatory ligands to CD28, and its upregulation on myeloid cells may be indicative of a pro-inflammatory anti-tumor response [42, 43]. Tim-3+ dendritic cells have been implicated in regulating immune response against tumors through multiple mechanisms [44–46]. Additionally, tracking proliferating myeloid cells through Ki-67 expression can be a useful biomarker for cancer immunotherapy targeting myeloid cells. In summary we provide a panel to comprehensively profile the immune cell landscape and their functionality in the mouse tumor and spleen. This is the first published 30-color full-spectrum cytometry OMIP panel for profiling the mouse immune system. We envision that this panel can be applied not only to other tissues in the context of syngeneic mouse tumor models, but also to non-tumor models, providing a valuable tool for a wide range of immunology studies.

## Similarity to other published OMIPs

This OMIP panel has similarities with OMIP-057, OMIP-076, OMIP-079 for characterization of T Cells [47–49]. The panel also overlaps with OMIP-031, which aims to characterize tumor-infiltrating T cells and the expression of immunologic checkpoint in the MC38 model [1]. It also overlaps with OMIP-032 and OMIP-061 to an extent with respect to the characterization of myeloid cells [2, 21]. The panel also overlaps with OMIP-054, which is a mass cytometry panel to characterize innate and adaptive immune cell infiltration in the brain, spleen, and bone marrow of an orthotopic murine glioblastoma model [10]. OMIP-054 include markers that are frequently observed in central nervous system immune cell infiltrates, while our panel is focused on markers that are frequently observed in subcutaneous syngeneic tumor’s immune cell infiltrates. Another distinction between OMIP-054 and our panel is in the dissociation digestive enzymes; OMIP-054 was tested with tissues dissociated with the Miltenyi’s Neural tissue dissociation enzymes while our panel was tested against the tumors dissociated with a mixture of collagenase type 4 and DNAse 1. Different dissociation enzyme recipes may have a different effect on sensitive protein epitopes. Our panel is the first high-dimensional spectral panel that was optimized to characterize dissociated mouse syngeneic tumors.

**Table 1.** Summary table for the application of this OMIP

|  |  |
| --- | --- |
| Purpose | High dimensional profiling of mouse immune cells to include subsets of T, B, NK, M-MDSC, G-MDSC, Dendritic Cells, and Macrophages. |
| Species | Mouse C57/Bl6 |
| Cell type | Single cell suspension from mouse spleen (mechanically processed), or single cell suspension from tumors (mechanically and enzymatically processed). |
| Cross references | OMIP-031, OMIP-032, OMIP-054, OMIP-057, OMIP-061, OMIP-076, OMIP-079 |

**Table 2.** Summary table for the antibodies in this panel

| Specificity | Vendor | Fluorochrome | Clone | Purpose |
| --- | --- | --- | --- | --- |
| CD45 | BD | BUV395 | 30-F11 | Pan White Blood Cells |
| Live/Dead Blue | Thermo Fisher | Live/Dead Blue (DAPI channel) | N/A | Viability |
| MHC-II (I-A/I-E) | BD | BUV496 | M5/114.15.2 | Dendritic Cells / Activated T Cells |
| TCR beta | BD | BUV563 | H57-597 | Pan alpha/beta T cells |
| CD8 | BD | BUV615 | 53-6.7 | CD8 T cells and Dendritic cell subset |
| NK1.1 | BD | BUV661 | PK136 | NK cells |
| PD-1 (CD279) | BD | BUV737 | J43 | T cell inhibitory receptor |
| CD103 | BD | BUV805 | M290 | Differentiation and Dendritic cell subset |
| Granzyme-B | Biolegend | BV421 | QA18A28 | NK cells & activated CD8 T cells |
| PDCA-1 (CD317) | Biolegend | Pacific Blue | 927 | Plasmacytoid DCs |
| PD-L1 (CD274) | BD | BV480 | MIH5 | Immunosuppressive myeloid cells |
| CD4 | Biolegend | BV510 | GK1.5 | CD4 T cells |
| CD11b | Biolegend | BV570 | M1/70 | Myeloid cells |
| Tim-3 (CD366) | Biolegend | BV605 | RMT3-23 | Co-inhibitory T & NK Cells |
| F4/80 | Biolegend | BV650 | BM8 | Macrophages |
| CD62L | Biolegend | BV711 | MEL-14 | Differentiation |
| CD80 | BD | BV750 | 16-10A1 | Activation |
| CD44 | BD | BV786 | IM7 | Memory T cells |
| Ki67 | BD | Alexa Fluor 488 | B56 | Proliferating cells |
| CD25 | BD | BB 700 | PC61 | Regulatory T cells |
| NKG2D | BD | PE | CX5 | NK Cells & Activated CD8 T Cells |
| Foxp3 | Thermo<br>Fisher | PE-eFluor610 | FJK-16s | Regulatory T cells |
| CD11c | Biolegend | PE-Cy5 | N418 | Dendritic Cells (DCs) |
| CD19 | Thermo<br>Fisher | PE-Cy5.5 | eBio1D3 | Pan B cells |
| LAG-3 (CD223) | Biolegend | PE-Cy7 | C9B7W | Exhausted T cells & NK cells |
| CD45R/B220 | Biolegend | PE-Fire810 | RA3-6B2 | B cells & plasmacytoid DCs |
| CTLA-4<br>(CD152) | Biolegend | APC | UC10-4B9 | Co-inhibitory |
| CD3 | Biolegend | Spark NIR 685 | 17A2 | Pan T cells |
| Ly6C | Biolegend | Alexa Fluor 700 | HK1.4 | Myeloid cells & M-MDSC |
| Ly6G | Biolegend | APC-Fire750 | 1A8 | Granulocytes & G-MDSC |

## Acknowledgments

We would like to thank Manfred Kraus, Kimberly Ly, Ofir Stefanson, Sunitha Bachawal, Priti Singh, Jennifer Sahni, Michael Sun, Olivia Perng, and Tanja Mehlo-Jensen for providing the mouse tissues, dissociation protocol, and critical inputs to optimize the panels.

## Conflict of Interest

Gabriel DeNiro, Kathryn Que, Jeong Kim, Bridget Schneider, Anandaroop Mukhopadhyay, and Anandi Sawant are employees of Bristol Myers Squibb. Tuan Andrew Nguyen and Soo Min Koo were employees of Bristol Myers Squibb. The authors declare no conflict of interest. The design, study conduct, and financial support for this research were provided by Bristol Myers Squibb. Bristol Myers Squibb participated in the interpretation of data, review, and approval of the publication. This study was performed with IACUC approval from Bristol Myers Squibb (ACUP# 21-041).

## Notes

### Competing Interest Statement

The authors have declared no competing interest.

## References

1. Nemoto, S., et al., OMIP-031: Immunologic checkpoint expression on murine effector and memory T-cell subsets. Cytometry A, 2016. 89(5): p. 427–9.

2. Unsworth, A., et al., OMIP-032: Two multi-color immunophenotyping panels for assessing the innate and adaptive immune cells in the mouse mammary gland. Cytometry A, 2016. 89(6): p. 527–30.

3. Waldman, A.D., J.M. Fritz, and M.J. Lenardo, A guide to cancer immunotherapy: from T cell basic science to clinical practice. Nat Rev Immunol, 2020. 20(11): p. 651–668.

4. Das, S. and D.B. Johnson, Immune-related adverse events and anti-tumor efficacy of immune checkpoint inhibitors. J Immunother Cancer, 2019. 7(1): p. 306.

5. Zhou, B., et al., Acquired Resistance to Immune Checkpoint Blockades: The Underlying Mechanisms and Potential Strategies. Front Immunol, 2021. 12: p. 693609.

6. Varayathu, H., et al., Combination Strategies to Augment Immune Check Point Inhibitors Efficacy - Implications for Translational Research. Front Oncol, 2021. 11: p. 559161.

7. Selby, M.J., et al., Preclinical Development of Ipilimumab and Nivolumab Combination Immunotherapy: Mouse Tumor Models, In Vitro Functional Studies, and Cynomolgus Macaque Toxicology. PLoS One, 2016. 11(9): p. e0161779.

8. Yu, J.W., et al., Tumor-immune profiling of murine syngeneic tumor models as a framework to guide mechanistic studies and predict therapy response in distinct tumor microenvironments. PLoS One, 2018. 13(11): p. e0206223.

9. Taylor, M.A., et al., Longitudinal immune characterization of syngeneic tumor models to enable model selection for immune oncology drug discovery. J Immunother Cancer, 2019. 7(1): p. 328.

10. Dusoswa, S.A., J. Verhoeff, and J.J. Garcia-Vallejo, OMIP-054: Broad Immune Phenotyping of Innate and Adaptive Leukocytes in the Brain, Spleen, and Bone Marrow of an Orthotopic Murine Glioblastoma Model by Mass Cytometry. Cytometry A, 2019. 95(4): p. 422–426.

11. Bendall, S.C., et al., A deep profiler’s guide to cytometry. Trends Immunol, 2012. 33(7): p. 323–32.

12. Hey, Y.Y., J.K. Tan, and H.C. O’Neill, Redefining Myeloid Cell Subsets in Murine Spleen. Front Immunol, 2015. 6: p. 652.

13. Rose, S., A. Misharin, and H. Perlman, A novel Ly6C/Ly6G-based strategy to analyze the mouse splenic myeloid compartment. Cytometry A, 2012. 81(4): p. 343–50.

14. Merad, M., et al., The dendritic cell lineage: ontogeny and function of dendritic cells and their subsets in the steady state and the inflamed setting. Annu Rev Immunol, 2013. 31: p. 563–604.

15. Omi, A., et al., Mature resting Ly6C(high) natural killer cells can be reactivated by IL-15. Eur J Immunol, 2014. 44(9): p. 2638–47.

16. Wensveen, F.M., V. Jelencic, and B. Polic, NKG2D: A Master Regulator of Immune Cell Responsiveness. Front Immunol, 2018. 9: p. 441.

17. Ghasemi, R., et al., Selective targeting of IL-2 to NKG2D bearing cells for improved immunotherapy. Nat Commun, 2016. 7: p. 12878.

18. Easom, N.J.W., et al., IL-15 Overcomes Hepatocellular Carcinoma-Induced NK Cell Dysfunction. Front Immunol, 2018. 9: p. 1009.

19. Merino, A., et al., Chronic stimulation drives human NK cell dysfunction and epigenetic reprograming. J Clin Invest, 2019. 129(9): p. 3770–3785.

20. Sordo-Bahamonde, C., et al., LAG-3 Blockade with Relatlimab (BMS-986016) Restores Anti-Leukemic Responses in Chronic Lymphocytic Leukemia. Cancers (Basel), 2021. 13(9).

21. DiPiazza, A.T., et al., OMIP-061: 20-Color Flow Cytometry Panel for High-Dimensional Characterization of Murine Antigen-Presenting Cells. Cytometry A, 2019. 95(12): p. 1226–1230.

22. Cabeza-Cabrerizo, M., et al., Dendritic Cells Revisited. Annu Rev Immunol, 2021. 39: p. 131–166.

23. Ghosn, E.E., et al., Distinct progenitors for B-1 and B-2 cells are present in adult mouse spleen. Proc Natl Acad Sci U S A, 2011. 108(7): p. 2879–84.

24. Largeot, A., et al., The B-side of Cancer Immunity: The Underrated Tune. Cells, 2019. 8(5).

25. Davis, M.M. and P.J. Bjorkman, T-cell antigen receptor genes and T-cell recognition. Nature, 1988. 334(6181): p. 395–402.

26. Fleming, C., et al., gammadelta T Cells: Unexpected Regulators of Cancer Development and Progression. Trends Cancer, 2017. 3(8): p. 561–570.

27. Liu, Q., Z. Sun, and L. Chen, Memory T cells: strategies for optimizing tumor immunotherapy. Protein Cell, 2020. 11(8): p. 549–564.

28. Larimer, B.M., et al., The Effectiveness of Checkpoint Inhibitor Combinations and Administration Timing Can Be Measured by Granzyme B PET Imaging. Clin Cancer Res, 2019. 25(4): p. 1196–1205.

29. Kim, K.H., et al., The First-week Proliferative Response of Peripheral Blood PD-1(+)CD8(+) T Cells Predicts the Response to Anti-PD-1 Therapy in Solid Tumors. Clin Cancer Res, 2019. 25(7): p. 2144–2154.

30. Kamphorst, A.O., et al., Proliferation of PD-1+ CD8 T cells in peripheral blood after PD-1-targeted therapy in lung cancer patients. Proc Natl Acad Sci U S A, 2017. 114(19): p. 4993–4998.

31. Banchereau, R., et al., Intratumoral CD103+ CD8+ T cells predict response to PD-L1 blockade. J Immunother Cancer, 2021. 9(4).

32. Miggelbrink, A.M., et al., CD4 T-Cell Exhaustion: Does It Exist and What Are Its Roles in Cancer? Clin Cancer Res, 2021. 27(21): p. 5742–5752.

33. Wherry, E.J. and M. Kurachi, Molecular and cellular insights into T cell exhaustion. Nat Rev Immunol, 2015. 15(8): p. 486–99.

34. Limagne, E., et al., Tim-3/galectin-9 pathway and mMDSC control primary and secondary resistances to PD-1 blockade in lung cancer patients. Oncoimmunology, 2019. 8(4): p. e1564505.

35. Huang, R.Y., et al., Compensatory upregulation of PD-1, LAG-3, and CTLA-4 limits the efficacy of single-agent checkpoint blockade in metastatic ovarian cancer. Oncoimmunology, 2017. 6(1): p. e1249561.

36. O’Rourke, K., Relatlimab plus nivolumab beneficial for previously untreated metastatic or unresectable melanoma. Cancer, 2022. 128(10): p. 1887.

37. Robert, C., et al., Nivolumab in previously untreated melanoma without BRAF mutation. N Engl J Med, 2015. 372(4): p. 320–30.

38. Larkin, J., F.S. Hodi, and J.D. Wolchok, Combined Nivolumab and Ipilimumab or Monotherapy in Untreated Melanoma. N Engl J Med, 2015. 373(13): p. 1270–1.

39. Pu, Y. and Q. Ji, Tumor-Associated Macrophages Regulate PD-1/PD-L1 Immunosuppression. Front Immunol, 2022. 13: p. 874589.

40. Tang, F. and P. Zheng, Tumor cells versus host immune cells: whose PD-L1 contributes to PD-1/PD-L1 blockade mediated cancer immunotherapy? Cell Biosci, 2018. 8: p. 34.

41. Lau, J., et al., Tumour and host cell PD-L1 is required to mediate suppression of anti-tumour immunity in mice. Nat Commun, 2017. 8: p. 14572.

42. Salomon, R. and R. Dahan, Next Generation CD40 Agonistic Antibodies for Cancer Immunotherapy. Front Immunol, 2022. 13: p. 940674.

43. Chen, X., et al., ILT4 inhibition prevents TAM- and dysfunctional T cell-mediated immunosuppression and enhances the efficacy of anti-PD-L1 therapy in NSCLC with EGFR activation. Theranostics, 2021. 11(7): p. 3392–3416.

44. de Mingo Pulido, A., et al., TIM-3 Regulates CD103(+) Dendritic Cell Function and Response to Chemotherapy in Breast Cancer. Cancer Cell, 2018. 33(1): p. 60–74 e6.

45. Chiba, S., et al., Tumor-infiltrating DCs suppress nucleic acid-mediated innate immune responses through interactions between the receptor TIM-3 and the alarmin HMGB1. Nat Immunol, 2012. 13(9): p. 832–42.

46. Wolf, Y., A.C. Anderson, and V.K. Kuchroo, TIM3 comes of age as an inhibitory receptor. Nat Rev Immunol, 2020. 20(3): p. 173–185.

47. Buus, T.B., M.H. Jee, and N. Odum, OMIP-057: Mouse gammadelta T-Cell Development Characterized by a 14 Color Flow Cytometry Panel. Cytometry A, 2019. 95(7): p. 726–729.

48. Mincham, K.T., J.D. Young, and D.H. Strickland, OMIP 076: High-dimensional immunophenotyping of murine T-cell, B-cell, and antibody secreting cell subsets. Cytometry A, 2021. 99(9): p. 888–892.

49. Natalini, A., et al., OMIP-079: Cell cycle of CD4(+) and CD8(+) naive/memory T cell subsets, and of Treg cells from mouse spleen. Cytometry A, 2021. 99(12): p. 1171–1175.

50. Mahnke, Y.D. and M. Roederer, Optimizing a multicolor immunophenotyping assay. Clin Lab Med, 2007. 27(3): p. 469–85, v.

51. Ferrer-Font, L., et al., Panel Optimization for High-Dimensional Immunophenotyping Assays Using Full-Spectrum Flow Cytometry. Curr Protoc, 2021. 1(9): p. e222.

52. DeNiro, G., P. Singh, and A. Nguyen, Performance comparison of the Spectral Signature, Spillover Spreading Matrix, and Data Unmixing Quality between the spectral Cytek Aurora and the spectral BD FACS Symphony A5 SE by a 35-Color Human Broad Immunophenotyping Panel. ScienceOpen Posters, 2022.

53. Sun, Y., et al., Robust Ki67 detection in human blood by flow cytometry for clinical studies. Bioanalysis, 2016. 8(23): p. 2399–2413.

