## Supplementary Material for "A 30-Color Full-Spectrum Flow Cytometry Panel to Characterize the Immune Cell Landscape in Spleen and Tumors within a Syngeneic MC-38 Murine Colon Carcinoma Model"

**Supplementary Figure 1. Instrument Configuration.** Below is the configuration of the FACS Symphony A5 SE with spectral unmixing capabilities.

| Laser | Detector | Mirror | Filter | Parameter | Corresponding Fluorochrome |
|---|---|---|---|---|---|
| UV 355nm 65mW | A | 765 LP | 809/32 BP | UV809 | BUV805 |
| | B | 704 LP | 736/64 BP | UV736 | BUV737 |
| | C | 675 LP | 695/40 BP | UV695 | |
| | D | 645 LP | 660/30 BP | UV660 | BUV661 |
| | E | 595 LP | 610/30 BP | UV610 | BUV615 |
| | F | 570 LP | 585/30 BP | UV585 | BUV563 |
| | G | 535 LP | 540/20 BP | UV540 | |
| | H | 495 LP | 515/60 BP | UV515 | BUV496 |
| | I | 425 LP | 446.5/67 BP | UV446 | Live/Dead Blue |
| | J | 365 LP | 379/34 BP | UV379 | BUV395 |
| Violet 405 nm 200mW | A | 810 LP | 845/70 BP | V810 | |
| | B | 765 LP | 785/50 BP | V785 | BV786 |
| | C | 730 LP | 750/40 BP | V750 | BV750 |
| | D | 690 LP | 710/40 BP | V710 | BV711 |
| | E | 665 LP | 680/30 BP | V680 | |
| | F | 645 LP | 660/30 BP | V660 | BV650 |
| | G | 605 LP | 615/25 BP | V615 | BV605 |
| | H | 585 LP | 595/30 BP | V595 | |
| | I | 570 LP | 576/20 BP | V576 | BV570 |
| | J | 530 LP | 540/20 BP | V540 | |
| | K | 495 LP | 510/40 BP | V510 | BV510 |
| | L | 465 LP | 470/15 BP | V470 | BV480 |
| | M | 430 LP | 450/40 BP | V450 | Pacific Blue |
| | N | 415 LP | 427/25 BP | V427 | BV421 |
| Blue 488 nm 150mW | A | 770 LP | 810/79 BP | B810 | |
| | B | 724 LP | 750/60 BP | B750 | |
| | C | 685 LP | 710/50 BP | B710 | BB700 |
| | D | 665 LP | 675/20 BP | B675 | |
| | E | 645 LP | 660/30 BP | B660 | |
| | F | 585 LP | 602/40 BP | B602 | |

| | | | | | |
|---|---|---|---|---|---|
| | G | 570 LP | 576/20 BP | B576 | |
| | H | 520 LP | 537/32 BP | B537 | Alexa Fluor 488 |
| | I | 500 LP | 510/20 BP | B510 | |
| | J | -- | 488/10 BP | SSC | |
| Yellow-Green 561 nm 150mW | A | 800 LP | 825.5/49 BP | YG825 | PE-Fire810 |
| | B | 750 LP | 780/60 BP | YG780 | PE-Cy7 |
| | C | 735 LP | 750/40 BP | YG750 | |
| | D | 699 LP | 730/50 BP | YG730 | |
| | E | 680 LP | 690/40 BP | YG690 | PE-Cy5.5 |
| | F | 665 LP | 670/20 BP | YG670 | |
| | G | 645 LP | 660/30 BP | YG660 | PE-Cy5 |
| | H | 595 LP | 602/40 BP | YG602 | PE-eFluor610 |
| | I | 570 LP | 585/30 BP | YG585 | PE |
| Red 637 nm 140mW | A | 750 LP | 780/60 BP | R780 | APC-Fire750 |
| | B | 720 LP | 730/50 BP | R730 | Alexa Fluor 700 |
| | C | 699 LP | 710/25 BP | R710 | |
| | D | 680 LP | 680/30 BP | R680 | Spark NIR 685 |
| | E | 665 LP | 675/20 BP | R675 | |
| | F | 645 LP | 660/30 BP | R660 | APC |

**Supplementary Figure 2.** Reagents with selected titer and manufacturer information.

| Specificity | Vendor | Fluorochrome | Clone | Catalog Number | Concentration (ug/ml) | Titer (ul/test) |
|---|---|---|---|---|---|---|
| Ki67 | BD | Alexa Fluor 488 | B56 | 558616 | 50 | 5 |
| CD25 | BD | BB700 | PC61 | 566498 | 200 | 1.25 |
| NKG2D | BD | PE | CX5 | 558403 | 200 | 1.25 |
| Foxp3 | Thermo Fisher | PE-eFluor610 | FJK-16s | 61-5773-82 | 200 | 1.25 |
| CD11c | Biolegend | PE-Cy5 | N418 | 117316 | 200 | 0.25 |
| CD19 | Thermo Fisher | PE-Cy5.5 | eBio1D3 | 35-0193-82 | 200 | 2.5 |
| LAG-3 (CD223) | Biolegend | PE-Cy7 | C9B7W | 125226 | 200 | 1.25 |
| CD45R/B220 | Biolegend | PE-Fire810 | RA3-6B2 | 103287 | 200 | 1.25 |
| Granzyme-B | Biolegend | BV421 | QA18A28 | 396414 | 100 | 5 |
| PDCA-1 (CD317) | Biolegend | Pacific Blue | 927 | 127018 | 500 | 0.5 |
| PD-L1 (CD274) | BD | BV480 | MIH5 | 746275 | 200 | 1.25 |
| CD4 | Biolegend | BV510 | GK1.5 | 100449 | 200 | 1.25 |
| CD11b | Biolegend | BV570 | M1/70 | 101233 | 100 | 1 |
| Tim-3 (CD366) | Biolegend | BV605 | RMT3-23 | 119721 | 200 | 1.25 |
| F4/80 | Biolegend | BV650 | BM8 | 123149 | 200 | 0.5 |
| CD62L | Biolegend | BV711 | MEL-14 | 104445 | 200 | 1.25 |
| CD80 | BD | BV750 | 16-10A1 | 747436 | 200 | 1.25 |
| CD44 | BD | BV786 | IM7 | 563736 | 200 | 0.15 |
| CD45 | BD | BUV395 | 30-F11 | 564279 | 200 | 0.5 |
| Live/Dead Blue | Thermo Fisher | DAPI | | L34962 | | 1:2000 |
| MHC-II (I-A/I-E) | BD | BUV496 | M5/114.15.2 | 750281 | 200 | 0.25 |
| TCR beta | BD | BUV563 | H57-597 | 748406 | 200 | 2.5 |
| CD8 | BD | BUV615 | 53-6.7 | 613004 | 200 | 1.25 |
| NK1.1 | BD | BUV661 | PK136 | 741477 | 200 | 1.25 |
| PD-1 (CD279) | BD | BUV737 | J43 | 749422 | 200 | 1.25 |

| | | | | | | |
|---|---|---|---|---|---|---|
| CD103 | BD | BUV805 | M290 | 741948 | 200 | 2.5 |
| CTLA-4 (CD152) | Biolegend | APC | UC10-4B9 | 106310 | 200 | 2 |
| CD3 | Biolegend | Spark NIR 685 | 17A2 | 100262 | 500 | 2 |
| Ly6C | Biolegend | Alexa Fluor 700 | HK1.4 | 128024 | 500 | 0.2 |
| Ly6G | Biolegend | APC-Fire750 | 1A8 | 127652 | 200 | 0.5 |

**Supplementary Figure 3. Panel Development.** Initially, we survey the literature and past data that was generated at Bristol Myer Squibb to propose a list of 40 markers that were frequently used and relevant in preclinical syngeneic tumor models. These markers can classify canonical immune cells and characterize the immune cells' activation/functionality. Due to commercial reagent availability and instrument configuration limitation, 29 markers along with 1 viability dye were selected in the final panel. Other markers that were proposed but later deprioritized in this panel were CD68, CD24, XCR1, CD172a, CD90.2, CD206, ICOS (CD278), PDPN, CD83, CD86, TIGIT**.** Good standard practices were implemented in designing this high dimensional flow cytometry panel [50, 51]. **A) Antigen classification**. Based on the literature as well as experience with these markers, antigens were organized into three groups: primary, secondary, and tertiary. Primary antigens are well-characterized antigens with clear expression pattern such as CD4, CD8, etc. Secondary antigens are well-characterized antigens with a continuum expression. Tertiary antigens are either antigens with low or uncharacterized expression, or critical markers that requires the best possible resolution based on the experiment's requirements. **B) Experiment iterations of panel development.** In the first iteration, the selection of fluorochrome and antigen was based on antigen classification and fluorochrome brightness on the BD FACS Symphony A5 SE, which was previously described [52]. We also consider spillover spread matrix (SSM) and antigen co-expression to maximize staining resolution. Blue boxes indicate reagents, which were changed at individual iterations. Markers, which were not changed after the first iteration are highlighted in green.

**S3A)**

| Antigen Classification | Markers |
|---|---|
| Primary | CD19, CD11c, CD45R/B220, PDCA-1, CD4, CD8, CD11b, F4/80, CD45, CD3, NK1.1, Ly6-C, Ly6-G, TCR-β, Live/Dead. |
| Secondary | CD25, PD-L1, CD62L, CD44, I/A-I/E (MHC-II), PD-1, CD103, CD80. |
| Tertiary | Ki67, NKG2D, LAG-3 (CD223), Granzyme B, Tim-3 (CD366), CTLA-4, Foxp3. |

**S3B)**

| Fluorochrome | Iteration 1 | Iteration 2 | Iteration 3 | Reasons |
|---|---|---|---|---|
| Alexa Fluor 488 | Ki67<br>(Clone Ki67) | Ki67<br>(Clone B56) | Ki67<br>(Clone B56) | Clone B56 was superior to clone Ki67. This was also previously reported in the literature [53]. |

| | | | | |
|---|---|---|---|---|
| NovaFluor Blue 610 | CD19 | CD19 | | The NovaFluor Blue 610 fluorochrome has non-specific binding with intratumoral myeloid cells even though the sample was blocked with Cellblox Blocking Buffer according to the manufacturer's recommendation. As a result, CD19 was moved from Novafluor Blue 610 to PE-Cy5.5. |
| BB700 | CD25 | CD25 | CD25 | |
| PE | NKG2D | NKG2D | NKG2D | |
| PE-eFluor610 | Foxp3 | Foxp3 | Foxp3 | |
| PE-Cy5 | CD11c | CD11c | CD11c | |
| PE-Cy5.5 | | | CD19 | CD19 was moved from NovaFluor Blue 610 to PE-Cy5.5. |
| PE-Cy7 | LAG-3 (CD223) | LAG-3 (CD223) | LAG-3 (CD223) | |
| PE-Fire810 | CD45R/B220 | CD45R/B220 | CD45R/B220 | |
| BV421 | Siglec H (Clone 440c) | Granzyme B (Clone QA18A28) | Granzyme B (Clone QA18A28) | Siglec H epitope was sensitive to the tumor dissociation enzyme per instructions. PDCA-1 was selected in subsequent iterations as an alternative marker to classify plasmacytoid dendritic cells. |
| Pacific Blue | Granzyme B (Clone GB11) | PDCA-1 (CD317) | PDCA-1 (CD317) | Granzyme B clone GB11 was not compatible with the eBioScience Foxp3 Fix/Perm buffer. Biolegend suggested clone QA18A28, which demonstrated excellent compatibility with the eBioScience Foxp3 Fix/Perm Buffer. |
| BV480 | PD-L1 (CD274) | PD-L1 (CD274) | PD-L1 (CD274) | |
| BV510 | CD24 | CD4 | CD4 | CD4 was prioritized over CD24 |
| Spark Violet 538 | CD4 | | | Spark Violet 538 was heavily impacted by BV510 due to similar spectral fingerprints. |
| BV570 | CD11b | CD11b | CD11b | |
| BV605 | Tim-3 (CD366) | Tim-3 (CD366) | Tim-3 (CD366) | |
| BV650 | F4/80 | F4/80 | F4/80 | |
| BV711 | CD62L | CD62L | CD62L | |

| | | | | |
|---|---|---|---|---|
| BV750 | CD80 | CD80 | CD80 | |
| BV786 | CD44 | CD44 | CD44 | |
| BUV395 | CD45 | CD45 | CD45 | |
| DAPI | Live/Dead Blue | Live/Dead Blue | Live/Dead Blue | |
| BUV496 | MHC-II (I-A/I-E) | MHC-II (I-A/I-E) | MHC-II (I-A/I-E) | |
| BUV563 | CD3 (Clone 145-2C11) | CD3 (Clone 17A2) | TCR Beta | CD3 (clone 17A2) was better than clone 145-2C11, but BUV563 was not a bright enough fluorochrome. Ultimately, CD3 (clone 17A2) in Spark NIR 685 was selected. TCR beta was later added to allow distinction between alpha/beta T cells and gamma/delta T cells. |
| BUV615 | CD8 | CD8 | CD8 | |
| BUV661 | NKp46 | NK1.1 | NK1.1 | Nkp46 staining was weaker than NK1.1 |
| BUV737 | PD-1 (CD279) | PD-1 (CD279) | PD-1 (CD279) | |
| BUV805 | CD103 | CD103 | CD103 | |
| APC | CTLA-4 (CD152) | CTLA-4 (CD152) | CTLA-4 (CD152) | |
| Spark NIR 685 | | | CD3 (Clone 17A2) | CD3 was moved from BUV563 to Spark NIR685 to improve the staining resolution. |
| Alexa Fluor 700 | Ly6C | Ly6C | Ly6C | |
| APC-Fire750 | Ly6G | Ly6G | Ly6G | |

**Supplementary Figure 4: (A)** Theoretical spill-over spreading error matrix (SSM) was obtained with Ultracomp compensation beads, Ultracomp compensation beads plus and in the case of Live/Dead Blue, Arc amine reactive compensation beads. The reagents from the final panel (iteration #3; supplementary table 3) were used. Values greater than or equal to 4 were highlighted in red, indicating that these combinations of fluorochromes need to be carefully assessed due to loss of resolution. **(B)** In this table, we describe the rationale for each challenging fluorochrome combination that has a Spread Error greater than or equal to 4. One strategy to overcome loss of resolution is to assign these fluorochromes to mutually exclusive antigens. For example, the matrix indicates that BUV661 can negatively impact the resolution of APC (Spread Error = 12.92); therefore, we assign BUV661 to NK1.1 (an NK-cell marker) and APC to CTLA-4 (a T-cell marker) to overcome the possible loss of resolution due to spread errors. In some cases where these fluorochromes cannot be assigned to mutually exclusive antigens, it's imperative to assess whether the loss of resolution is acceptable. For example, the matrix suggests that BUV496 can negatively impact the resolution of BUV395 (SE = 7.61); 2x2 gating analysis confirms excellent CD45 BUV395 staining despite possible BUV496 spread error. Alternatively, we can prioritize and accept logical compromise based on *a priori* understanding of the biological models and the drug mechanism of action. For example, the matrix suggests that PE-Cy5.5 can negatively impact the resolution of BB700 (SE = 8.90). We decided to assign PE-Cy5.5 and BB700 mutually exclusively gating-antigens (PE-Cy5.5: CD19 which is restricted to B cells, BB700: CD25 which is used to identify T-regulatory cells). While it is also possible for some activated B cells to express CD25, our understanding of the MC38 tumor model suggests that this biological phenomenon can be deprioritized.

**S4A)**

| Spread Donor (Fluorochrome) \ Spread Receiver (Detector) | BUV395 | Live/Dead Blue | BUV496 | BUV563 | BUV615 | BUV661 | BUV737 | BUV805 | BV421 | PacBlue | BV480 | BV510 | BV570 | BV605 | BV650 | BV711 | BV750 | BV786 | Alexa Fluor 488 | BB700 | PE | PE-eFluor610 | PE-Cy5 | PE-Cy5-5 | PE-Cy7 | PE-Fire810 | APC | Spark NIR 685 | Alexa Fluor 700 | APC-Fire750 |
|---|---|---|---|---|---|---|---|---|---|---|---|---|---|---|---|---|---|---|---|---|---|---|---|---|---|---|---|---|---|---|
| BUV395 | 0.00 | 2.72 | 0.63 | 0.39 | 0.26 | 0.12 | 0.07 | 0.11 | 0.12 | 0.00 | 0.00 | 0.00 | 0.20 | 0.22 | 0.13 | 0.08 | 0.11 | 0.16 | 0.00 | 0.08 | 0.26 | 0.16 | 0.09 | 0.00 | 0.00 | 0.00 | 0.12 | 0.00 | 0.08 | 0.00 |
| Live/Dead Blue | 2.40 | 0.00 | 2.04 | 0.87 | 0.78 | 0.27 | 0.00 | 0.09 | 3.08 | 4.67 | 1.25 | 0.73 | 0.18 | 0.22 | 0.15 | 0.09 | 0.08 | 0.07 | 0.06 | 0.12 | 0.21 | 0.42 | 0.15 | 0.00 | 0.00 | 0.06 | 0.48 | 0.00 | 0.10 | 0.11 |
| BUV496 | 7.61 | 2.81 | 0.00 | 2.03 | 1.75 | 0.81 | 0.36 | 0.24 | 0.17 | 0.60 | 1.64 | 1.60 | 1.01 | 0.79 | 0.44 | 0.22 | 0.16 | 0.10 | 0.60 | 0.15 | 0.58 | 0.78 | 0.16 | 0.09 | 0.00 | 0.00 | 0.50 | 0.17 | 0.17 | 0.00 |
| BUV563 | 1.35 | 0.56 | 0.66 | 0.00 | 3.27 | 1.24 | 0.53 | 0.29 | 0.15 | 0.10 | 0.72 | 0.37 | 0.91 | 0.70 | 0.37 | 0.24 | 0.14 | 0.08 | 0.79 | 0.38 | 1.87 | 1.97 | 0.52 | 0.37 | 0.23 | 0.18 | 0.80 | 0.39 | 0.29 | 0.00 |
| BUV615 | 0.49 | 0.32 | 0.17 | 1.30 | 0.00 | 1.94 | 1.06 | 0.57 | 0.09 | 0.09 | 0.16 | 0.12 | 0.40 | 0.71 | 0.64 | 0.53 | 0.34 | 0.19 | 0.12 | 0.75 | 1.04 | 3.31 | 0.98 | 0.79 | 0.49 | 0.34 | 1.31 | 0.71 | 0.57 | 0.12 |
| BUV661 | 2.69 | 1.27 | 0.26 | 0.43 | 1.05 | 0.00 | 2.01 | 1.00 | 0.12 | 0.17 | 0.41 | 0.09 | 0.21 | 0.24 | 1.93 | 0.89 | 0.72 | 0.40 | 0.03 | 0.70 | 0.18 | 0.54 | 1.39 | 0.84 | 0.63 | 0.41 | 12.92 | 6.24 | 2.51 | 1.22 |
| BUV737 | 1.83 | 0.87 | 0.28 | 0.22 | 0.24 | 0.31 | 0.00 | 2.36 | 0.12 | 0.00 | 0.00 | 0.00 | 0.09 | 0.12 | 0.21 | 0.94 | 1.76 | 0.67 | 0.03 | 1.58 | 0.12 | 0.10 | 0.28 | 0.71 | 0.58 | 0.36 | 0.39 | 0.97 | 7.29 | 1.37 |
| BUV805 | 2.22 | 1.41 | 0.44 | 0.39 | 0.38 | 0.20 | 0.53 | 0.00 | 0.14 | 0.00 | 0.37 | 0.09 | 0.23 | 0.17 | 0.13 | 0.07 | 0.40 | 0.78 | 0.00 | 0.11 | 0.16 | 0.15 | 0.00 | 0.06 | 0.49 | 0.50 | 0.11 | 0.27 | 0.36 | 1.49 |
| BV421 | 0.31 | 1.50 | 0.44 | 0.34 | 0.26 | 0.10 | 0.07 | 0.05 | 0.00 | 3.70 | 2.96 | 1.43 | 0.60 | 0.35 | 0.20 | 0.11 | 0.07 | 0.08 | 0.05 | 0.06 | 0.15 | 0.22 | 0.00 | 0.00 | 0.07 | 0.12 | 0.00 | 0.05 | 0.05 | 0.00 |
| PacBlue | 0.10 | 0.31 | 0.43 | 0.42 | 0.35 | 0.17 | 0.12 | 0.05 | 0.65 | 0.00 | 4.44 | 2.31 | 1.01 | 0.68 | 0.39 | 0.19 | 0.15 | 0.11 | 0.11 | 0.06 | 0.28 | 0.26 | 0.11 | 0.00 | 0.00 | 0.00 | 0.14 | 0.14 | 0.06 | 0.08 |
| BV480 | 0.34 | 0.74 | 1.30 | 0.98 | 0.87 | 0.36 | 0.21 | 0.11 | 0.59 | 10.55 | 0.00 | 4.41 | 2.05 | 1.42 | 0.80 | 0.42 | 0.29 | 0.17 | 0.51 | 0.11 | 0.51 | 0.46 | 0.12 | 0.09 | 0.03 | 0.00 | 0.20 | 0.11 | 0.10 | 0.00 |
| BV510 | 0.27 | 0.22 | 1.39 | 2.08 | 1.96 | 0.97 | 0.57 | 0.37 | 0.42 | 0.72 | 2.71 | 0.00 | 2.89 | 2.37 | 1.72 | 0.91 | 0.76 | 0.48 | 0.10 | 0.18 | 0.57 | 0.70 | 0.22 | 0.08 | 0.00 | 0.00 | 0.51 | 0.30 | 0.27 | 0.20 |
| BV570 | 0.17 | 0.22 | 0.29 | 1.96 | 1.72 | 0.88 | 0.53 | 0.27 | 1.12 | 1.11 | 1.46 | 0.86 | 0.00 | 2.54 | 1.69 | 0.89 | 0.67 | 0.38 | 0.15 | 0.39 | 1.37 | 1.62 | 0.55 | 0.44 | 0.33 | 0.25 | 0.47 | 0.37 | 0.22 | 0.10 |
| BV605 | 0.21 | 0.17 | 0.16 | 1.33 | 3.55 | 1.64 | 0.97 | 0.47 | 1.05 | 0.84 | 0.85 | 0.26 | 1.08 | 0.00 | 2.91 | 1.54 | 1.18 | 0.71 | 0.14 | 0.68 | 1.03 | 2.06 | 0.87 | 0.74 | 0.51 | 0.40 | 0.98 | 0.53 | 0.48 | 0.11 |
| BV650 | 0.18 | 0.15 | 0.10 | 0.25 | 0.92 | 2.95 | 1.35 | 0.65 | 0.80 | 0.76 | 0.71 | 0.27 | 0.28 | 0.73 | 0.00 | 2.20 | 1.46 | 0.85 | 0.00 | 0.48 | 0.14 | 0.44 | 0.70 | 0.58 | 0.41 | 0.27 | 3.00 | 2.44 | 1.19 | 0.56 |
| BV711 | 0.26 | 0.21 | 0.12 | 0.16 | 0.14 | 0.63 | 3.92 | 1.37 | 1.00 | 1.05 | 0.99 | 0.34 | 0.21 | 0.20 | 0.63 | 0.00 | 2.80 | 1.85 | 0.00 | 0.82 | 0.13 | 0.22 | 0.26 | 0.58 | 0.50 | 0.33 | 0.72 | 2.00 | 3.43 | 1.19 |
| BV750 | 0.15 | 0.17 | 0.21 | 0.37 | 0.29 | 0.16 | 1.83 | 1.18 | 3.15 | 1.82 | 1.85 | 0.99 | 0.63 | 0.58 | 0.43 | 0.95 | 0.00 | 2.07 | 0.06 | 0.36 | 0.30 | 0.45 | 0.17 | 0.22 | 0.33 | 0.27 | 0.27 | 0.35 | 1.01 | 0.69 |
| BV786 | 0.47 | 0.36 | 0.17 | 0.19 | 0.15 | 0.15 | 1.12 | 2.63 | 3.06 | 2.21 | 1.22 | 0.54 | 0.29 | 0.23 | 0.22 | 0.41 | 2.60 | 0.00 | 0.06 | 0.14 | 0.18 | 0.02 | 0.08 | 0.09 | 0.42 | 0.37 | 0.10 | 0.24 | 0.55 | 1.12 |
| Alexa Fluor 488 | 0.00 | 0.00 | 0.28 | 0.35 | 0.35 | 0.18 | 0.08 | 0.08 | 0.17 | 0.00 | 0.81 | 0.38 | 0.59 | 0.37 | 0.24 | 0.19 | 0.08 | 0.08 | 0.00 | 0.24 | 0.90 | 1.34 | 0.29 | 0.18 | 0.06 | 0.00 | 0.10 | 0.19 | 0.00 | 0.00 |
| BB700 | 0.00 | 0.00 | 1.07 | 1.88 | 1.68 | 1.22 | 1.61 | 1.16 | 0.87 | 0.96 | 3.48 | 2.19 | 2.63 | 2.09 | 1.95 | 3.49 | 2.19 | 1.51 | 0.74 | 0.00 | 1.52 | 2.44 | 2.05 | 1.45 | 1.00 | 0.79 | 2.38 | 4.27 | 2.25 | 1.10 |
| PE | 0.13 | 0.00 | 0.21 | 1.95 | 1.37 | 0.56 | 0.29 | 0.13 | 0.33 | 0.33 | 0.79 | 0.60 | 2.47 | 1.63 | 0.89 | 0.59 | 0.23 | 0.15 | 0.59 | 0.80 | 0.00 | 3.61 | 0.99 | 0.64 | 0.40 | 0.25 | 0.44 | 0.40 | 0.16 | 0.00 |
| PE-eFluor610 | 0.05 | 0.00 | 0.11 | 0.58 | 1.70 | 0.58 | 0.29 | 0.12 | 0.09 | 0.27 | 0.51 | 0.47 | 0.89 | 0.87 | 0.81 | 0.81 | 0.28 | 0.16 | 0.40 | 1.23 | 1.61 | 0.00 | 1.20 | 0.77 | 0.53 | 0.33 | 0.56 | 0.42 | 0.20 | 0.07 |
| PE-Cy5 | 0.07 | 0.00 | 0.06 | 0.25 | 0.46 | 1.32 | 0.75 | 0.40 | 0.18 | 0.00 | 0.00 | 0.00 | 0.32 | 0.27 | 1.39 | 2.20 | 0.86 | 0.49 | 0.11 | 4.47 | 0.62 | 0.99 | 0.00 | 2.60 | 1.64 | 1.08 | 4.01 | 4.83 | 1.72 | 0.91 |
| PE-Cy5-5 | 0.11 | 0.08 | 0.05 | 0.31 | 0.30 | 0.55 | 1.39 | 0.60 | 0.31 | 0.00 | 0.24 | 0.26 | 0.47 | 0.35 | 1.32 | 3.46 | 1.17 | 0.72 | 0.19 | 8.90 | 1.19 | 1.13 | 2.73 | 0.00 | 2.71 | 1.44 | 1.22 | 4.73 | 1.52 | 0.72 |
| PE-Cy7 | 0.17 | 0.08 | 0.06 | 0.13 | 0.11 | 0.08 | 0.33 | 0.98 | 0.12 | 0.00 | 0.14 | 0.00 | 0.18 | 0.15 | 0.10 | 0.31 | 0.77 | 1.21 | 0.03 | 0.57 | 0.40 | 0.38 | 0.20 | 0.48 | 0.00 | 2.97 | 0.11 | 0.21 | 0.30 | 1.16 |
| PE-Fire810 | 0.15 | 0.05 | 0.07 | 0.38 | 0.35 | 0.17 | 0.22 | 0.82 | 0.12 | 0.00 | 0.06 | 0.06 | 0.57 | 0.43 | 0.26 | 0.49 | 0.47 | 0.94 | 0.14 | 0.88 | 1.25 | 1.27 | 0.43 | 0.71 | 2.10 | 0.00 | 0.35 | 0.69 | 0.30 | 0.59 |
| APC | 0.09 | 0.06 | 0.12 | 0.16 | 0.14 | 1.50 | 0.84 | 0.44 | 0.11 | 0.00 | 0.63 | 0.00 | 0.09 | 0.23 | 1.82 | 0.90 | 0.77 | 0.40 | 0.00 | 0.83 | 0.00 | 0.51 | 1.67 | 1.08 | 0.82 | 0.57 | 0.00 | 5.68 | 2.72 | 1.49 |
| Spark NIR 685 | 0.10 | 0.00 | 0.09 | 0.17 | 0.17 | 0.44 | 0.50 | 0.33 | 0.08 | 0.00 | 0.00 | 0.00 | 0.15 | 0.21 | 0.51 | 0.74 | 0.57 | 0.35 | 0.00 | 0.47 | 0.17 | 0.36 | 0.71 | 0.69 | 0.67 | 0.42 | 2.92 | 0.00 | 2.57 | 1.55 |
| Alexa Fluor 700 | 0.08 | 0.00 | 0.08 | 0.15 | 0.12 | 0.19 | 0.95 | 0.54 | 0.15 | 0.00 | 0.39 | 0.00 | 0.18 | 0.17 | 0.22 | 1.37 | 1.57 | 0.82 | 0.00 | 0.71 | 0.16 | 0.23 | 0.26 | 1.17 | 0.86 | 0.58 | 0.55 | 1.77 | 0.00 | 2.67 |
| APC-Fire750 | 0.13 | 0.05 | 0.00 | 0.07 | 0.11 | 0.17 | 0.30 | 1.03 | 0.12 | 0.00 | 0.55 | 0.00 | 0.29 | 0.11 | 0.20 | 0.17 | 0.70 | 1.44 | 0.00 | 0.14 | 0.25 | 0.11 | 0.19 | 0.17 | 1.52 | 1.53 | 0.64 | 0.57 | 0.73 | 0.00 |

**S4B)**

| Theoretical Spread Error greater than 4.0 | Strategy to overcome the possible loss of resolution |
|---|---|
| BUV661 spread into APC (SE = 12.92) | BUV661 and APC were assigned to mutually exclusively gating-antigens (BUV661: NK1.1 which is restricted to NK cells, APC: CTLA-4 which is restricted to T cells) |
| BV480 spread into Pacific Blue | 2x2 gating analysis confirms excellent PDCA-1 Pacific Blue staining despite possible BV480 |

| | |
|---|---|
| (SE = 10.55) | spread error. |
| PE-Cy5.5 spread into BB700 (SE = 8.90) | PE-Cy5.5 and BB700 were assigned to mutually exclusively gating-antigens (PE-Cy5.5: CD19 which is restricted to B cells, BB700: CD25 which is used to identify T-regulatory cells). While it is also possible for activated B cells to express CD25, it is outside of the scope of this work. |
| BUV496 spread into BUV395 (SE = 7.61) | 2x2 gating analysis confirms excellent CD45 BUV395 staining despite possible BUV496 spread error. |
| BUV737 spread into Alexa Fluor 700 (SE = 7.29) | BUV737 and Alexa Fluor 700 were assigned to mutually exclusively gating-antigens (BUV737: PD-1 which is typically expressed on T cells, Alexa Fluor 700: Ly6c which is typically highly expressed on myeloid cells). While it is also possible for some T cells to express Ly6c, it is outside of the scope of this work. |
| BUV661 spread into Spark NIR 685 (SE = 6.24) | BUV661 and Spark NIR 685 were assigned to mutually exclusively gating antigens (BUV661: NK1.1 which is typically expressed on NK cells, Spark NIR 685: CD3 which is expressed on T cells). It's known that NK-T cells can express both NK1.1 and CD3; 2x2 gating analysis confirms excellent resolution of NK-T cells. |
| APC spread into Spark NIR 685 (SE = 5.68) | 2x2 gating analysis confirms excellent Spark NIR 685 CD3 staining despite possible spread from APC CTLA-4. |
| PE-Cy5 spread into Spark NIR 685 (SE = 4.83) | PE-Cy5 and Spark NIR 685 were assigned to mutually exclusively gating antigens (PE-Cy5: CD11c which is typically expressed on myeloid cells, Spark NIR 685: CD3 which is typically expressed on T cells) |
| PE-Cy5.5 spread into Spark NIR 685 (SE = 4.73) | PE-Cy5.5 and Spark NIR 685 were assigned to mutually exclusively gating antigens (PE-Cy5.5: CD19 which is typically expressed on B cells, Spark NIR 685: CD3 which is typically expressed on T cells) |
| Live/Dead Blue spread into Pacific Blue (SE = 4.67) | 2x2 gating analysis confirms excellent PDCA-1 Pacific Blue staining despite possible spread error from Live/Dead Blue. |
| PE-Cy5 spread into BB700 (SE = | PE-Cy5 and BB700 were assigned to mutually exclusively gating antigens (PE-Cy5: CD11c |

| | |
|---|---|
| 4.47) | which is typically expressed on myeloid cells, BB700: CD25 which is typically expressed on T cells) |
| Pacific Blue spread into BV480 (SE = 4.44) | Pacific Blue was assigned to PDCA-1, which is typically expressed on plasmacytoid dendritic cells. BV480 was assigned to PD-L1, and in the context of MC-38 tumor, PD-L1 is highly expressed on tumor-associated macrophages, MDSC, conventional dendritic cells, and to a much lesser extent, plasmacytoid dendritic cells. |
| BV480 spread into BV510 (SE = 4.41) | BV480 and BV510 were assigned to mutually exclusively gating antigens (BV480: PD-L1 which is typically expressed on myeloid cells and tumor cells, BV510: CD4 which is typically expressed on T cells) |
| BB700 spread into Spark NIR 685 (SE = 4.27) | 2x2 gating analysis confirms excellent Spark NIR 685 CD3 staining despite possible BB700 CD25 spread error. |
| PE-Cy5 spread into APC (SE = 4.01) | PE-Cy5 and APC were assigned to mutually exclusively gating antigens (PE-Cy5: CD11c which is typically expressed on myeloid cells, APC: CTLA-4 which is typically expressed on T cells) |

**Supplementary Figure 5: (A)** Normalized spectral signatures of reagents in the panel. **(B)** Similarity Matrix of the Reagents in the Panel. Overall, the fluorochrome reagents have very distinct spectral signatures from each other, with similarity index ranging from 0 to 0.83. This indicates that the reagents can be easily unmixed from each other.

**S5A)**

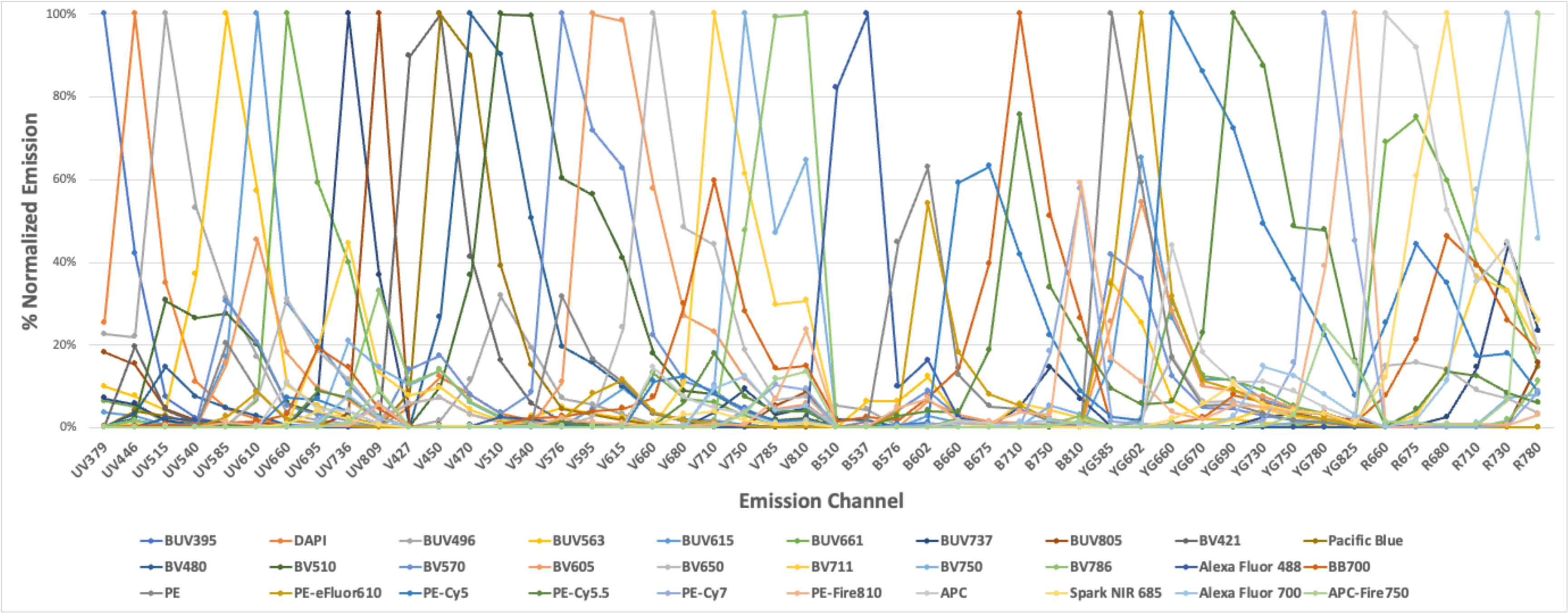

## S5B)

| | BUV395 | DAPI | BUV496 | BUV563 | BUV615 | BUV661 | BUV737 | BUV805 | BV421 | Pacific Blue | BV480 | BV510 | BV570 | BV605 | BV650 | BV711 | BV750 | BV786 | Alexa Fluor 488 | BB700 | PE | PE-eFluor610 | PE-Cy5 | PE-Cy5.5 | PE-Cy7 | PE-Fire810 | APC | Spark NIR 685 | Alexa Fluor 700 | APC-Fire750 |
|---|---|---|---|---|---|---|---|---|---|---|---|---|---|---|---|---|---|---|---|---|---|---|---|---|---|---|---|---|---|---|
| BUV395 | 1.00 | 0.58 | 0.29 | 0.11 | 0.04 | 0.04 | 0.07 | 0.22 | 0.06 | 0.02 | 0.03 | 0.03 | 0.01 | 0.01 | 0.01 | 0.00 | 0.01 | 0.01 | 0.00 | 0.00 | 0.00 | 0.00 | 0.00 | 0.00 | 0.00 | 0.00 | 0.00 | 0.00 | 0.00 | 0.00 |
| DAPI | 0.58 | 1.00 | 0.51 | 0.15 | 0.05 | 0.04 | 0.06 | 0.19 | 0.27 | 0.17 | 0.15 | 0.14 | 0.05 | 0.03 | 0.02 | 0.02 | 0.03 | 0.03 | 0.01 | 0.01 | 0.01 | 0.00 | 0.00 | 0.00 | 0.00 | 0.00 | 0.00 | 0.00 | 0.00 | 0.00 |
| BUV496 | 0.29 | 0.51 | 1.00 | 0.42 | 0.16 | 0.05 | 0.04 | 0.11 | 0.12 | 0.18 | 0.38 | 0.52 | 0.16 | 0.11 | 0.04 | 0.02 | 0.02 | 0.02 | 0.08 | 0.02 | 0.07 | 0.03 | 0.01 | 0.01 | 0.00 | 0.01 | 0.01 | 0.00 | 0.00 | 0.00 |
| BUV563 | 0.11 | 0.15 | 0.42 | 1.00 | 0.61 | 0.10 | 0.03 | 0.05 | 0.02 | 0.02 | 0.07 | 0.25 | 0.38 | 0.33 | 0.07 | 0.02 | 0.01 | 0.01 | 0.07 | 0.04 | 0.47 | 0.34 | 0.06 | 0.06 | 0.01 | 0.08 | 0.03 | 0.01 | 0.01 | 0.00 |
| BUV615 | 0.04 | 0.05 | 0.16 | 0.61 | 1.00 | 0.28 | 0.09 | 0.04 | 0.01 | 0.01 | 0.03 | 0.15 | 0.35 | 0.52 | 0.21 | 0.07 | 0.03 | 0.02 | 0.01 | 0.07 | 0.40 | 0.57 | 0.22 | 0.15 | 0.03 | 0.10 | 0.11 | 0.03 | 0.02 | 0.02 |
| BUV661 | 0.04 | 0.04 | 0.05 | 0.10 | 0.28 | 1.00 | 0.34 | 0.08 | 0.01 | 0.00 | 0.01 | 0.05 | 0.07 | 0.17 | 0.45 | 0.30 | 0.08 | 0.04 | 0.00 | 0.36 | 0.05 | 0.08 | 0.43 | 0.19 | 0.03 | 0.04 | 0.77 | 0.63 | 0.33 | 0.12 |
| BUV737 | 0.07 | 0.06 | 0.04 | 0.03 | 0.09 | 0.34 | 1.00 | 0.38 | 0.01 | 0.00 | 0.01 | 0.02 | 0.02 | 0.05 | 0.15 | 0.50 | 0.29 | 0.20 | 0.00 | 0.31 | 0.01 | 0.01 | 0.12 | 0.16 | 0.11 | 0.08 | 0.18 | 0.22 | 0.49 | 0.26 |
| BUV805 | 0.22 | 0.19 | 0.11 | 0.05 | 0.04 | 0.08 | 0.38 | 1.00 | 0.03 | 0.01 | 0.02 | 0.03 | 0.02 | 0.02 | 0.04 | 0.15 | 0.18 | 0.31 | 0.00 | 0.07 | 0.01 | 0.01 | 0.02 | 0.03 | 0.09 | 0.09 | 0.03 | 0.04 | 0.09 | 0.25 |
| BV421 | 0.06 | 0.27 | 0.12 | 0.02 | 0.01 | 0.01 | 0.01 | 0.03 | 1.00 | 0.76 | 0.41 | 0.20 | 0.17 | 0.06 | 0.08 | 0.09 | 0.14 | 0.12 | 0.01 | 0.00 | 0.01 | 0.00 | 0.00 | 0.00 | 0.00 | 0.00 | 0.00 | 0.00 | 0.00 | 0.00 |
| Pacific Blue | 0.02 | 0.17 | 0.18 | 0.02 | 0.01 | 0.00 | 0.00 | 0.01 | 0.76 | 1.00 | 0.76 | 0.40 | 0.16 | 0.07 | 0.07 | 0.08 | 0.12 | 0.10 | 0.01 | 0.00 | 0.01 | 0.01 | 0.00 | 0.00 | 0.00 | 0.00 | 0.00 | 0.00 | 0.00 | 0.00 |
| BV480 | 0.03 | 0.15 | 0.38 | 0.07 | 0.03 | 0.01 | 0.01 | 0.02 | 0.41 | 0.76 | 1.00 | 0.78 | 0.25 | 0.15 | 0.07 | 0.05 | 0.07 | 0.06 | 0.04 | 0.02 | 0.06 | 0.02 | 0.01 | 0.00 | 0.00 | 0.01 | 0.01 | 0.01 | 0.00 | 0.01 |
| BV510 | 0.03 | 0.14 | 0.52 | 0.25 | 0.15 | 0.05 | 0.02 | 0.03 | 0.20 | 0.40 | 0.78 | 1.00 | 0.56 | 0.42 | 0.19 | 0.08 | 0.07 | 0.05 | 0.03 | 0.07 | 0.16 | 0.06 | 0.02 | 0.02 | 0.01 | 0.02 | 0.02 | 0.01 | 0.01 | 0.01 |
| BV570 | 0.01 | 0.05 | 0.16 | 0.38 | 0.35 | 0.07 | 0.02 | 0.02 | 0.17 | 0.16 | 0.25 | 0.56 | 1.00 | 0.76 | 0.28 | 0.08 | 0.06 | 0.05 | 0.02 | 0.10 | 0.57 | 0.40 | 0.10 | 0.08 | 0.02 | 0.10 | 0.05 | 0.02 | 0.01 | 0.01 |
| BV605 | 0.01 | 0.03 | 0.11 | 0.33 | 0.52 | 0.17 | 0.05 | 0.02 | 0.06 | 0.07 | 0.15 | 0.42 | 0.76 | 1.00 | 0.54 | 0.17 | 0.11 | 0.08 | 0.01 | 0.17 | 0.41 | 0.47 | 0.19 | 0.13 | 0.04 | 0.09 | 0.11 | 0.03 | 0.03 | 0.02 |
| BV650 | 0.01 | 0.02 | 0.04 | 0.07 | 0.21 | 0.45 | 0.15 | 0.04 | 0.08 | 0.07 | 0.07 | 0.19 | 0.28 | 0.54 | 1.00 | 0.44 | 0.21 | 0.14 | 0.00 | 0.37 | 0.08 | 0.11 | 0.28 | 0.15 | 0.03 | 0.04 | 0.35 | 0.22 | 0.15 | 0.06 |
| BV711 | 0.00 | 0.02 | 0.02 | 0.02 | 0.07 | 0.30 | 0.50 | 0.15 | 0.09 | 0.08 | 0.05 | 0.08 | 0.08 | 0.17 | 0.44 | 1.00 | 0.64 | 0.46 | 0.00 | 0.59 | 0.01 | 0.02 | 0.15 | 0.22 | 0.09 | 0.09 | 0.22 | 0.30 | 0.48 | 0.22 |
| BV750 | 0.01 | 0.03 | 0.02 | 0.01 | 0.03 | 0.08 | 0.29 | 0.18 | 0.14 | 0.12 | 0.07 | 0.07 | 0.06 | 0.11 | 0.21 | 0.64 | 1.00 | 0.83 | 0.00 | 0.29 | 0.01 | 0.01 | 0.05 | 0.09 | 0.13 | 0.14 | 0.05 | 0.05 | 0.19 | 0.20 |
| BV786 | 0.01 | 0.03 | 0.02 | 0.01 | 0.02 | 0.04 | 0.20 | 0.31 | 0.12 | 0.10 | 0.06 | 0.05 | 0.05 | 0.08 | 0.14 | 0.46 | 0.83 | 1.00 | 0.00 | 0.20 | 0.01 | 0.01 | 0.03 | 0.06 | 0.15 | 0.19 | 0.03 | 0.03 | 0.12 | 0.26 |
| Alexa Fluor 488 | 0.00 | 0.01 | 0.08 | 0.07 | 0.01 | 0.00 | 0.00 | 0.00 | 0.01 | 0.01 | 0.04 | 0.03 | 0.02 | 0.01 | 0.00 | 0.00 | 0.00 | 0.00 | 1.00 | 0.03 | 0.09 | 0.06 | 0.01 | 0.01 | 0.00 | 0.01 | 0.00 | 0.00 | 0.00 | 0.00 |
| BB700 | 0.00 | 0.01 | 0.02 | 0.04 | 0.07 | 0.36 | 0.31 | 0.07 | 0.00 | 0.00 | 0.02 | 0.07 | 0.10 | 0.17 | 0.37 | 0.59 | 0.29 | 0.20 | 0.03 | 1.00 | 0.08 | 0.10 | 0.47 | 0.54 | 0.17 | 0.16 | 0.34 | 0.45 | 0.38 | 0.17 |
| PE | 0.00 | 0.01 | 0.07 | 0.47 | 0.40 | 0.05 | 0.01 | 0.01 | 0.01 | 0.01 | 0.06 | 0.16 | 0.57 | 0.41 | 0.08 | 0.01 | 0.01 | 0.01 | 0.09 | 0.08 | 1.00 | 0.77 | 0.15 | 0.13 | 0.03 | 0.18 | 0.05 | 0.01 | 0.01 | 0.01 |
| PE-eFluor610 | 0.00 | 0.00 | 0.03 | 0.34 | 0.57 | 0.08 | 0.01 | 0.01 | 0.00 | 0.01 | 0.02 | 0.06 | 0.40 | 0.47 | 0.11 | 0.02 | 0.01 | 0.01 | 0.06 | 0.10 | 0.77 | 1.00 | 0.29 | 0.18 | 0.04 | 0.16 | 0.09 | 0.02 | 0.01 | 0.01 |
| PE-Cy5 | 0.00 | 0.00 | 0.01 | 0.06 | 0.22 | 0.43 | 0.12 | 0.02 | 0.00 | 0.00 | 0.01 | 0.02 | 0.10 | 0.19 | 0.28 | 0.15 | 0.05 | 0.03 | 0.01 | 0.47 | 0.15 | 0.29 | 1.00 | 0.66 | 0.17 | 0.16 | 0.54 | 0.37 | 0.21 | 0.10 |
| PE-Cy5.5 | 0.00 | 0.00 | 0.01 | 0.06 | 0.15 | 0.19 | 0.16 | 0.03 | 0.00 | 0.00 | 0.00 | 0.02 | 0.08 | 0.13 | 0.15 | 0.22 | 0.09 | 0.06 | 0.01 | 0.54 | 0.13 | 0.18 | 0.66 | 1.00 | 0.37 | 0.30 | 0.20 | 0.21 | 0.22 | 0.13 |
| PE-Cy7 | 0.00 | 0.00 | 0.00 | 0.01 | 0.03 | 0.03 | 0.11 | 0.09 | 0.00 | 0.00 | 0.00 | 0.01 | 0.02 | 0.04 | 0.03 | 0.09 | 0.13 | 0.15 | 0.00 | 0.17 | 0.03 | 0.04 | 0.17 | 0.37 | 1.00 | 0.76 | 0.05 | 0.04 | 0.12 | 0.33 |
| PE-Fire810 | 0.00 | 0.00 | 0.01 | 0.08 | 0.10 | 0.04 | 0.08 | 0.09 | 0.00 | 0.00 | 0.01 | 0.02 | 0.10 | 0.09 | 0.04 | 0.09 | 0.14 | 0.19 | 0.01 | 0.16 | 0.18 | 0.16 | 0.16 | 0.30 | 0.76 | 1.00 | 0.05 | 0.04 | 0.08 | 0.25 |
| APC | 0.00 | 0.00 | 0.01 | 0.03 | 0.11 | 0.77 | 0.18 | 0.03 | 0.00 | 0.00 | 0.01 | 0.02 | 0.05 | 0.11 | 0.35 | 0.22 | 0.05 | 0.03 | 0.00 | 0.34 | 0.05 | 0.09 | 0.54 | 0.20 | 0.05 | 0.05 | 1.00 | 0.72 | 0.41 | 0.15 |
| Spark NIR 685 | 0.00 | 0.00 | 0.00 | 0.01 | 0.03 | 0.63 | 0.22 | 0.04 | 0.00 | 0.00 | 0.01 | 0.01 | 0.02 | 0.03 | 0.22 | 0.30 | 0.05 | 0.03 | 0.00 | 0.45 | 0.01 | 0.02 | 0.37 | 0.21 | 0.04 | 0.04 | 0.72 | 1.00 | 0.53 | 0.22 |
| Alexa Fluor 700 | 0.00 | 0.00 | 0.00 | 0.01 | 0.02 | 0.33 | 0.49 | 0.09 | 0.00 | 0.00 | 0.00 | 0.01 | 0.01 | 0.03 | 0.15 | 0.48 | 0.19 | 0.12 | 0.00 | 0.38 | 0.01 | 0.01 | 0.21 | 0.22 | 0.12 | 0.08 | 0.41 | 0.53 | 1.00 | 0.43 |
| APC-Fire750 | 0.00 | 0.00 | 0.00 | 0.00 | 0.02 | 0.12 | 0.26 | 0.25 | 0.00 | 0.00 | 0.01 | 0.01 | 0.01 | 0.02 | 0.06 | 0.22 | 0.20 | 0.26 | 0.00 | 0.17 | 0.01 | 0.01 | 0.10 | 0.13 | 0.33 | 0.25 | 0.15 | 0.22 | 0.43 | 1.00 |

**Supplementary Figure 6: Tissue Processing Protocols**

**Required Materials:**

| **Reagent** | **source** | **Catalog** |
|---|---|---|
| Collagenase type 4 | Worhtington | LS004188 |
| DNAse 1 | Sigma Aldrich | 1010459001 |
| Calcium chloride 1M | VWR | AAJ63122-AD |
| GentleMACS C Tubes | Miltenyi Biotec | 130-093-237 |
| HBSS without Calcium and Magnesium | Corning | 21-021-CM |
| RMPI-1640 | Corning | 10-040-CV |
| FBS (heat inactivated) | Gibco | 10082147 |
| EDTA (0.5M) | Invitrogen | 15575020 |
| 100um Cell strainer | Corning | 352360 |
| 50ml Conical Tube | Corning | 352070 |
| GentleMACS Octo Dissociator with Heaters | Miltenyi Biotec | 130-096-427 |

| **Digest Mix (3mls of Digest per 1 gram of Tumor)** |
|---|
| 250U/ml Collagense type 4 |
| 100ug/ml DNAse1 |
| 5mM CaCl2 |
| 5% FBS |
| in HBSS |
| **(Keep at room temperature)** |

| **Quench (Keep it cold on ice)** |
|---|
| 10% FBS |
| 2mM EDTA |
| in RPMI 1640 |

| R10 Media |
|---|
| 50mL heat-inactivated fetal bovine serum |
| 450mL RPMI-1640 media |

| Complete media (Keep it cold on ice) |
|---|
| 419.5mL RPMI-1640 media |
| 50mL heat-inactivated fetal bovine serum |
| 5mL (5e4 I.U. / 5e4ug) Penicillin / Streptomycin (Cellgro, #30-002-C1) |
| 5mL (2.38 x 10-2 M) Sodium Bicarbonate (Sigma, #S-8761) |
| 5mL (1x) Non-Essential Amino Acids (Cellgro, #25-025-C1) |
| 5mL (2 x 10-3 M) L-glutamine (Cellgro, #25-005-C1) |
| 5mL (1 x 10-2 M) HEPES buffer (Gibco, #15630-080) |
| 5mL (1 x 10-3 M) Sodium Pyruvate (Sigma, #S8636) |
| 0.5mL (5 x 10-5 M) Beta-Mercaptoethanol (Sigma, #M7522) |
| Sterile filter into 500mL bottle |

**Tumor Processing Protocols**

1. Transfer tumors to C tubes
2. Chop tumors with scissors in C tubes 20-30 times to break into a 2-4mm small chunks.
3. Add **3mls of Digest Mix** to each C tube, and close tightly (2 clicks)
4. Place C tubes in GentleMACS Octo Dissociator and run **m_tumor_imp_03 TWICE**
5. Place C tube on a rotating platform in a **37'C incubator for 30min** or a 37C shaking water bath for 30 mins
6. Remove C tubes from incubation and **vortex on high for 10secs**
7. Quench digest enzymes with serum and EDTA containing media by adding **7mls of Cold Quench Buffer**
8. Pour contents (~10mls) over a 100um filter into a labelled 50ml conical tubes
9. Scrub the filter with the back of syringe plunger to break up any remaining chunks

10 Rinse the C tube with another **5mls of Quench Buffer** and pour over filter (~15mls)
11 Spin down all tumors at 4'C 500xg for 5min and resuspend in **1ml Complete media**
12 Count at **1:10 Counting Dilution** for tumor-infiltrating cells on NucleoCounter
13 Bring cell volume to 2e7/ml

**Spleen processing- without enzymes**

1 Transfer spleens from 24 well plate to C tubes
2 Add 3mls of R10 media to each C tube, and close tightly (2 clicks)
3 Place C tubes in GentleMACS Octo Dissociator and run **m_spleen_03 TWICE**
4 Pour contents over a 100um filter into a labelled 50ml conical
5 Scrub the filter with the back of syringe plunger to break up any remaining chunks
6 Rinse the C tube with 7mls of R10 media and pour over filter
10 Spin down at 4'C 500xg for 5min
11 Resuspend in 3ml RBC lysis buffer and incubate for 5 minutes
12 Quench reaction with 7ml R10 media
13 Pour contents over a 100um filter into a Labelled 50ml conical
14 Rinse the C tube with 5mls of R10 and pour over filter
15 Spin down all spleens at 4'C 500xg for 5min and resuspend in **5ml Complete media**
16 Count at **1:20 Counting Dilution** on NucleoCounter
17 Bring cell volume to 2e7/ml

**Supplementary Figure 7**: **Flow Cytometry Staining Protocol (Surface + Intracellular Staining)**

**Required Materials:**

- Live/Dead Fixable Blue Dead Cell Staining Kit (Thermo Fisher, L34962)
- TruStain FcX (anti-mouse CD16/32) Antibody (Biolegend, 101320)
- TruStain Monocyte Blocker (Biolegend, 426103)
- FACS Stain Buffer (Thermo Fisher, 00-4222-26)
- Brilliant Stain Buffer Plus (BD Biosciences, 566385)
- eBioscience Foxp3 / Transcription Factor Staining Buffer Set (Thermo Fisher, 00-5523-00)
- Flow Cytometry antibody: spin the antibody vial at 10,000 x g immediately prior to making antibody cocktail. Make cocktail in the presence of Brilliant Stain Buffer Plus. Cocktails with BUV dyes are stable for 2 hours. Cocktails without BUV dyes are stable for 24 hours
- Ultracomp Beads (Thermo Fisher, 01-2222-42)

**Staining Protocol:**

1. Plate 1x10^7 cells per tube
2. Spin the plate at 350 x g for 5min; flick off supernatant
3. Resuspend cells in 200ul PBS
4. Spin the plate at 350 x g for 5min; flick off supernatant
5. Make Live/Dead Blue viability dye mix
   a) Reconstitute one vial of reactive dye with 50ul of DMSO
   b) Add 1ul of reconstituted dye to 5ml PBS

6. Add 50ul of viability dye mix (prepared in step 5b) to each tube (from step 4)
7. Incubate for 20 min at RT in dark
8. Add 150ul FACS Stain Buffer
9. Spin tubes at 500 x g for 5min; flick off supernatant
10. Make surface Fc block working solution

- For every 1ml needed, add 100ul Trustain FcX block and 100ul Trustain monocyte blocker (Biolegend) in 800ul FACS stain buffer

11. Add 50ul Fc block working solution (prepared in step 10) to each tube (from step 9).
12. Incubate 5 min at RT
13. Add 50ul surface stain cocktail (recipe described below)

### Surface Cocktail Recipe (per test)

| Specificity | Vendor | Fluorochrome | Clone | Catalog Number | Concentration (ug/ml) | Titer (ul/test) |
|---|---|---|---|---|---|---|
| CD25 | BD | BB700 | PC61 | 566498 | 200 | 1.25 |
| NKG2D | BD | PE | CX5 | 558403 | 200 | 1.25 |
| CD11c | Biolegend | PE-Cy5 | N418 | 117316 | 200 | 0.25 |
| CD19 | Thermo Fisher | PE-Cy5.5 | eBio1D3 | 35-0193-82 | 200 | 2.5 |
| LAG-3 (CD223) | Biolegend | PE-Cy7 | C9B7W | 125226 | 200 | 1.25 |
| CD45R/B220 | Biolegend | PE-Fire810 | RA3-6B2 | 103287 | 200 | 1.25 |
| PDCA-1 (CD317) | Biolegend | Pacific Blue | 927 | 127018 | 500 | 0.5 |
| PD-L1 (CD274) | BD | BV480 | MIH5 | 746275 | 200 | 1.25 |
| CD4 | Biolegend | BV510 | GK1.5 | 100449 | 200 | 1.25 |
| CD11b | Biolegend | BV570 | M1/70 | 101233 | 100 | 1 |
| Tim-3 (CD366) | Biolegend | BV605 | RMT3-23 | 119721 | 200 | 1.25 |
| F4/80 | Biolegend | BV650 | BM8 | 123149 | 200 | 0.5 |
| CD62L | Biolegend | BV711 | MEL-14 | 104445 | 200 | 1.25 |
| CD80 | BD | BV750 | 16-10A1 | 747436 | 200 | 1.25 |
| CD44 | BD | BV786 | IM7 | 563736 | 200 | 0.15 |
| CD45 | BD | BUV395 | 30-F11 | 564279 | 200 | 0.5 |
| MHC-II (I-A/I-E) | BD | BUV496 | M5/114.15.2 | 750281 | 200 | 0.25 |
| TCR beta | BD | BUV563 | H57-597 | 748406 | 200 | 2.5 |
| CD8 | BD | BUV615 | 53-6.7 | 613004 | 200 | 1.25 |
| NK1.1 | BD | BUV661 | PK136 | 741477 | 200 | 1.25 |
| PD-1 (CD279) | BD | BUV737 | J43 | 749422 | 200 | 1.25 |
| CD103 | BD | BUV805 | M290 | 741948 | 200 | 2.5 |
| CD3 | Biolegend | Spark NIR 685 | 17A2 | 100262 | 500 | 2 |
| Ly6C | Biolegend | Alexa Fluor 700 | HK1.4 | 128024 | 500 | 0.2 |

| | | | | | | |
|---|---|---|---|---|---|---|
| Ly6G | Biolegend | APC-Fire750 | 1A8 | 127652 | 200 | 0.5 |
| Brilliant Stain Buffer Plus | BD | | | 566385 | | 10 |
| FACS Stain Buffer | Thermo Fisher | | | 00-4222-26 | | 11.65 |
| | | | | | **Total** | **50** |

14. Pipette samples 3x to fully mix the cells
15. Incubate for 20 min at room temperature
16. Add 150ul FACS stain buffer
17. Spin the plate at 350 x g for 5min; flick off supernatant
18. Repeat step 17 1 time
19. Make 1x eBio fix/perm buffer
    - For every 1ml needed, add 250ul of 4x fix/perm concentrate in 750ul fix/perm diluent
20. Resuspend cells in 100ul 1X eBio fix/perm buffer
21. Pipette samples 3x to fully mix the cells
22. Incubate 30 min at RT in dark (Mouse samples can be incubated for up to 18 hours at 2-8°C in the dark)
23. Make 1x eBio Permeabilization buffer (need 1ml of each well)
    - For every 1ml needed, add 100ul of 10x Permeabilization buffer in 900ul distilled water
24. Add 100ul 1X Permeabilization Buffer
25. Spin the plate at 600 x g for 5min; flick off supernatant
26. Resuspend in 200ul 1X Permeabilization buffer
27. Spin the plate at 600 x g for 5min; flick off supernatant
28. Make intracellular Fc block working solution
    - For every 1ml needed, add 100ul Trustain FcX block, 100ul Trustain monocyte blocker in 800ul 1X Permeabilization Buffer.
29. Resuspend cells in 50ul intracellular Fc block working solution. Pipettes samples 3x. Incubate for 5 minutes
30. Add 50ul of Intracellular stain cocktail (recipe described below)

### Intracellular Cocktail Recipe (per test)

| Specificity | Vendor | Fluorochrome | Clone | Catalog Number | Concentration (ug/ml) | Titer (ul/test) |
|---|---|---|---|---|---|---|
| Ki67 | BD | Alexa Fluor 488 | B56 | 558616 | 50 | 5 |
| Foxp3 | Thermo Fisher | PE-eFluor610 | FJK-16s | 61-5773-82 | 200 | 1.25 |
| Granzyme-B | Biolegend | BV421 | QA18A28 | 396414 | 100 | 5 |

| | | | | | | |
|---|---|---|---|---|---|---|
| CTLA-4 (CD152) | Biolegend | APC | UC10-4B9 | 106310 | 200 | 2 |
| Brilliant Stain Buffer Plus | BD | | | 566385 | | 10 |
| 1X diluted Permeabilization Buffer | Thermo Fisher | | | 00-5523-00 | | 26.75 |
| | | | | | **Total** | **50** |

31. Pipette samples 3x to fully mix the cells
32. Incubate 45min at RT in dark
33. Add 150ul 1x Permeabilization Buffer
34. Spin the plate at 600 x g for 5min; flick off supernatant
35. Repeat wash 2 more time with 200ul 1X Permeabilization Buffer
36. Make 0.5% PFA
    - PFA Stock is 4%
    - For every 1ml needed, add 125ul 4% PFA in 875ul FACS Stain buffer
37. Resuspend cells in 250ul 0.5% PFA (prepared in step 36).

**Preparation of Single-Color Compensation Controls (Fluorochrome-conjugated antibody)**

1. Label a tube for each fluorochrome that will be used in the experiment.
2. Mix beads by pulse-vortexing.
3. Add 2 drops of UltraComp eBeads™ to each tube. Add 200ul of PBS to each tube.
4. Add 0.5ul of antibody conjugate to each tube.
5. Mix well by pulse-vortexing.
6. Incubate at room temperature for 5 minutes in the dark.
7. Add 2 mL of Flow Cytometry Staining Buffer to each tube and centrifuge at 400 x g for 5 minutes.
8. Decant supernatant and add 0.2mL of 0.5% PFA buffer to each tube.
9. Mix briefly by pulse-vortexing before analysis.

**Preparation of Single-Color Compensation Controls (Live/dead Dye with Arc Bead)**

1. Prepare 2 tubes; one tube with the Arc negative bead; one tube with the Arc positive bead.
2. Mix beads by pulse-vortexing.

3. Add 2 drops of ArcPositive bead to the positive tube; and 2 drops of ArcNegative Bead to the negative tube. Add 200ul of PBS to each tube.
4. Add 1ul of diluted live/dead dye to the ArcPositive tube.
5. Mix well by flicking, inverting vigorously, or pulse-vortexing.
6. Incubate at room temperature for 10 minutes in the dark.
7. Add 2 mL of Flow Cytometry Staining Buffer to each tube and centrifuge at 400 x g for 5 minutes.
8. Decant supernatant and add 0.2mL of 0.5% PFA buffer to each tube.
9. Mix briefly by pulse-vortexing before analysis.

| Specificity | Fluorochrome | Compensation Bead | Vendor | Catalog Number | Titer (ul/test) |
|---|---|---|---|---|---|
| Ki67 | Alexa Fluor 488 | Ultracomp eBeads | Thermo Fisher | 01-2222-42 | 1 |
| CD25 | BB700 | Ultracomp eBeads | Thermo Fisher | 01-2222-42 | 0.25 |
| NKG2D | PE | Ultracomp eBeads | Thermo Fisher | 01-2222-42 | 0.25 |
| Foxp3 | PE-eFluor610 | Ultracomp eBeads | Thermo Fisher | 01-2222-42 | 0.25 |
| CD11c | PE-Cy5 | Ultracomp eBeads | Thermo Fisher | 01-2222-42 | 0.25 |
| CD19 | PE-Cy5.5 | Ultracomp eBeads | Thermo Fisher | 01-2222-42 | 0.25 |
| CD223 (LAG-3) | PE-Cy7 | Ultracomp eBeads | Thermo Fisher | 01-2222-42 | 0.25 |
| CD45R/B220 | PE-Fire810 | Ultracomp eBeads | Thermo Fisher | 01-2222-42 | 0.25 |
| Granzyme-B | BV421 | Ultracomp eBeads | Thermo Fisher | 01-2222-42 | 0.25 |
| CD317 (PDCA-1) | Pacific Blue | Ultracomp eBeads | Thermo Fisher | 01-2222-42 | 1 |

| | | | | | |
|---|---|---|---|---|---|
| CD274 (PD-L1) | BV480 | Ultracomp eBeads | Thermo Fisher | 01-2222-42 | 1 |
| CD4 | BV510 | Ultracomp eBeads | Thermo Fisher | 01-2222-42 | 1 |
| CD11b | BV570 | Ultracomp eBeads | Thermo Fisher | 01-2222-42 | 1 |
| CD366 (Tim-3) | BV605 | Ultracomp eBeads | Thermo Fisher | 01-2222-42 | 1 |
| F4/80 | BV650 | Ultracomp eBeads | Thermo Fisher | 01-2222-42 | 1 |
| CD62L | BV711 | Ultracomp eBeads | Thermo Fisher | 01-2222-42 | 1 |
| CD80 | BV750 | Ultracomp eBeads | Thermo Fisher | 01-2222-42 | 1 |
| CD44 | BV786 | Ultracomp eBeads Plus | Thermo Fisher | 01-3333-42 | 1 |
| CD45 | BUV395 | Ultracomp eBeads | Thermo Fisher | 01-2222-42 | 1 |
| Live/Dead Blue | DAPI | Arc Amine Reactive Compensation Beads | Thermo Fisher | A10346 | 1:2000 |
| I-A/I-E (MHC-II) | BUV496 | Ultracomp eBeads | Thermo Fisher | 01-2222-42 | 1 |
| TCR beta | BUV563 | Ultracomp eBeads | Thermo Fisher | 01-2222-42 | 1 |
| CD8 | BUV615 | Ultracomp eBeads | Thermo Fisher | 01-2222-42 | 1 |
| NK1.1 | BUV661 | Ultracomp eBeads | Thermo Fisher | 01-2222-42 | 1 |
| CD279 (PD-1) | BUV737 | Ultracomp eBeads | Thermo Fisher | 01-2222-42 | 1 |

| | | | | | |
|---|---|---|---|---|---|
| CD103 | BUV805 | Ultracomp eBeads | Thermo Fisher | 01-2222-42 | 1 |
| CD152 (CTLA-4) | APC | Ultracomp eBeads | Thermo Fisher | 01-2222-42 | 1 |
| CD3 | Spark NIR 685 | Ultracomp eBeads | Thermo Fisher | 01-2222-42 | 1 |
| Ly6C | Alexa Fluor 700 | Ultracomp eBeads | Thermo Fisher | 01-2222-42 | 1 |
| Ly6G | APC-Fire750 | Ultracomp eBeads | Thermo Fisher | 01-2222-42 | 1 |
